# Continuous Rod Noise and Nonlinear Synaptic Filtering Shape Signal Detection in the Primary Rod Pathway

**DOI:** 10.64898/2026.09.29.754402

**Authors:** Ulisse Bocchero, Johan Pahlberg

**Affiliations:** Photoreceptor Physiology Group, National Eye Institute, National Institutes of Health, Bethesda, Maryland, 20892

**Keywords:** PDE6, single-photon detection, retinal noise, rod bipolar cells, nonlinear signal processing

## Abstract

Rod photoreceptors are optimized to detect single photons, but their performance near absolute visual threshold is limited by intrinsic phototransduction noise. The two major components are discrete single-photon-like events and continuous fluctuations in outer-segment current. Although spontaneous rhodopsin activation is established as the source of discrete noise, the molecular origin of continuous noise remains controversial, and how these distinct noise sources influence signal processing at the first visual synapse is not fully understood.

To address these questions, we combined genetic manipulation of rhodopsin, phosphodiesterase 6 (PDE6), and glutamic acid rich protein 2 (GARP2) with single-cell recordings from rods and rod-bipolar cells (RBCs) and pharmacological manipulation of mGluR6 signaling. Continuous noise was unaffected by reduced rhodopsin concentration but was strongly influenced by PDE6 expression and regulation, supporting basal PDE6 activity as a major source of continuous noise. Reducing either discrete or continuous noise increased signal-to-noise ratio (SNR) in rods, but these changes were differentially processed at the rod-to-RBC synapse. In WT retina, the nonlinear transformation increased SNR, whereas elevated continuous noise limited this increase. Conversely, reduced continuous noise increased rod SNR, which was maintained in RBCs and associated with a lower detection threshold. Pharmacological reduction of the mGluR6-dependent nonlinearity altered these genotype-dependent differences, indicating that nonlinear filtering is well matched to normal signal and noise distributions but does not fully compensate when presynaptic noise is experimentally altered.

Together, our study identifies spontaneous PDE6 activity as a major source of continuous noise and demonstrates that continuous noise interacts with nonlinear processing at the first visual synapse to determine the fidelity and detection threshold of dim-light signals.

**Significance Statement:** Vision near absolute threshold requires retinal circuits to distinguish single-photon responses from intrinsic photoreceptor noise. We show that basal PDE6 activity is a major source of continuous noise in rod photoreceptors and that altering this noise affects signal fidelity after convergence onto rod bipolar cells. Nonlinear filtering at the first visual synapse interacts with presynaptic noise levels, while reducing continuous rod noise improves the detection of near-threshold light signals. These findings link the molecular mechanisms generating photoreceptor noise to its processing by downstream retinal circuitry and demonstrate how noise control at both cellular and synaptic levels contributes to the remarkable sensitivity of the primary rod pathway.

## Introduction

The remarkable sensitivity of vision under dim-light conditions approaches the physical limit of photon detection (Hecht et al., 1942). In vertebrates, this exceptional sensitivity is achieved by rod photoreceptors, which are capable of reliably detecting and transmitting the absorption of a single photon (Baylor et al., 1979). Phototransduction begins when a photon of light photoisomerizes the visual pigment rhodopsin (Rh) within the rod outer segment. Activated rhodopsin catalyzes the activation of the heterotrimeric G protein transducin (Tr), which in turn activates cGMP phosphodiesterase-6 (PDE6). PDE6 then hydrolyzes cyclic guanosine monophosphate (cGMP), reducing the probability of opening cGMP-gated ion channels and producing the hyperpolarizing electrical light response of rods. The high gain and reproducibility of this signaling cascade enable rods to generate reliable responses to single photons. However, spontaneous activity generated in darkness produces intrinsic noise that ultimately limits the detection of weak light signals (Barlow, 1956, 1957; Baylor et al., 1984; Donner, 1992). Two major forms of such intrinsic dark noise have been identified in rod photoreceptors, called discrete and continuous noise (Baylor et al., 1980). Discrete noise consists of spontaneous current deflections whose amplitude and kinetics closely resemble genuine single-photon responses (Baylor et al., 1979) and arises from spontaneous activation of the visual pigment (Baylor et al., 1980; Bocchero and Pahlberg, 2023). Continuous noise consists of ongoing low-amplitude fluctuations in outer-segment currents and has traditionally been thought to arise downstream of visual pigment activation, with basal PDE6 activity considered a likely source (Lamb, 1987; Rieke and Baylor, 1996; Pahlberg and Sampath, 2011; Bocchero and Pahlberg 2023). However, recent evidence has suggested that visual pigment itself may contribute to continuous noise (Chai et al., 2024). Thus, although the origin of discrete noise is well established, the molecular source of continuous noise remains unresolved.

Resolving the origin of continuous noise is important for understanding how photoreceptors achieve reliable single-photon detection and how these signals are transmitted through retinal circuitry despite intrinsic noise. At the first visual synapse, each rod bipolar cell (RBC) receives convergent input from multiple rods. In mouse retina, approximately 20–25 rods contact an individual RBC (Tsukamoto et al., 2001; Berntson et al., 2004), forming the first stage of the highly sensitive primary rod pathway (Dacheux and Raviola, 1986; Smith et al., 1986). This convergence requires the rod-to-RBC synapse to distinguish between sparse photon responses arising in only a fraction of the rods from intrinsic photoreceptor noise originating in all of the rod population (Baylor et al., 1984; van Rossum and Smith, 1998). This discrimination is facilitated by a specialized metabotropic glutamate receptor 6 (mGluR6) -dependent signaling cascade in RBCs, which generates a strong nonlinear threshold at the rod-to-RBC synapse (Field and Rieke, 2002; Sampath and Rieke, 2004; Robson et al., 2004). This nonlinearity improves signal-to-noise ratio (SNR) by suppressing weak rod inputs and a substantial fraction of the continuous noise, while transmitting larger single-photon responses (Field and Rieke, 2002; Trexler et al., 2011). However, how different forms and levels of rod noise influence nonlinear filtering, and whether the threshold is optimized for altered signal-to-noise conditions, remain poorly understood.

To determine the origin of continuous noise and how intrinsic noise properties influence signal processing in the retinal circuitry, we combined complementary genetic, physiological, and pharmacological approaches. Using transgenic mouse models with altered expression of rhodopsin, PDE6, and the PDE6 regulatory protein GARP2, together with single-cell patch-clamp recordings from rods and rod bipolar cells and pharmacological manipulation of mGluR6-dependent signaling, we examined how different noise sources are processed at the first visual synapse. Our results support basal PDE6 activity as a major source of continuous dark noise in rod photoreceptors, and demonstrate that nonlinear processing at the first visual synapse interacts with continuous rod noise to determine the fidelity and detection threshold of weak light signals.

## Materials and Methods

### Mouse lines and preparation

This study was conducted in accordance with the recommendations of the *Guide for the Care and Use of Laboratory Animals* of the National Institutes of Health (NIH), and the Association for Research in Vision and Ophthalmology Statement for the Use of Animals in Ophthalmic and Vision Research. The animal use protocol was approved by the Animal Care and Use Committees of the NIH (approval ASP 1344-21).

All mice used in this study were 2–6 months of age, and approximately equal numbers of both sexes were used for experiments. Wild-type (WT) C57BL/6J mice were obtained from The Jackson Laboratory (Bar Harbor, ME, USA). Genetically modified mouse lines included Rh heterozygous (Rh^+/−^, Calvert et al., 2001), cGMP PDE6 heterozygous (PDE6^+/−^, Majumder et al., 2015) and GARP2 knockout (GARP2^−/−^, Stacks et al., 2026). All lines were bred for at least five generations onto a common C57BL/6J background before experimental use. As previously shown, the retinas of these animals remain structurally preserved at the ages examined and exhibit robust rod-driven bipolar-cell responses, consistent with largely preserved rod phototransduction and synaptic transmission (Lem et al., 1999; Majumder et al., 2015; Bocchero and Pahlberg, 2023; Stacks et al., 2026).

Animals were maintained under a 12-h light/12-h dark cycle in standard rodent housing with standard chow and were always dark adapted overnight before experimentation. All subsequent experimental manipulations were performed in darkness under infrared illumination (λmax ∼940 nm) and visualized using infrared image converters (B.E. Meyers). Mice were euthanized according to guidelines approved by NIH (ASP 1344–21). Following euthanasia, eyes were enucleated, the cornea and lens were removed, and eyecups were maintained in darkness at 32°C in bicarbonate-buffered Ames’ medium (Sigma-Aldrich, catalog #A1420) and equilibrated with 5% CO₂/95% O₂.

### Retinal slice preparation

Experiments were performed in the morning or early afternoon, and all experimental preparations were performed in darkness, under infrared illumination (λmax ∼940 nm, Thorlabs). The retina was carefully removed from the retinal pigmented epithelium (RPE) with fine forceps. Isolated retinal tissue was embedded in 3% low-gelling temperature agarose (Sigma-Aldrich, catalog #A-0701) in HEPES buffered Ames’ medium. Cross-sectional retinal slices (200 μm in thickness) were cut in chilled, oxygenated HEPES-buffered Ames’ medium using a vibrating microtome (VT-1000S; Leica). Slices were transferred either to a recording chamber or to a storage container for use later in the experiment. During recordings, the retinal slice was stabilized with a custom-made stainless-steel anchor, and consistently superfused with bicarbonate-buffered Ames’ medium at ∼8 ml/min. For experiments examining the rod-to-RBC nonlinearity, the mGluR6 antagonist LY341495 (Tocris Bioscience, catalog #1209) was added to the internal solution at a concentration of 0.4 µM. Bath temperature was maintained at 35 ± 1 °C using a temperature controller (TC-324B; Warner Instruments, Holliston, MA).

### Physiological recordings of rod photoreceptors and rod bipolar cells

Single-cell patch-clamp recordings were performed using a custom electrophysiology setup in the Photoreceptor Physiology Group at the National Institutes of Health. Physiological responses were recorded from individual rod photoreceptors and rod bipolar cells (RBCs) in retinal slices using previously established methods (Pahlberg et al., 2018; Bocchero and Pahlberg, 2023).

Rods and RBCs were visualized using a 40x, water immersion objective (Olympus) under infrared illumination provided by an LED source (λmax ∼940 nm; Thorlabs) attached to the transmitted-light path of the microscope (Zeiss). Rod somas were identified by morphology and location in the outer nuclear layer (ONL). RBC somas were identified by their characteristic morphology and location adjacent to the outer plexiform layer (OPL).

Patch pipettes were fabricated on the day of each experiment from filamented borosilicate-glass capillaries (BF120-69-10; Sutter Instruments, Novato, CA) using a vertical puller (Narishige), to a tip resistance in the bath medium of 12-15 MΩ. The standard pipette internal solution for all whole-cell patch-clamp recordings contained (in mM): 125 K-aspartate, 10 KCl, 10 HEPES, 5 N-methyl glucamine-HEDTA, 0.5 CaCl_2_, 1 ATP-Mg, and 0.2 GTP-Mg; pH was adjusted to 7.3 with N-methyl glucamine-OH and osmolarity to 280 mOsm.

All the experiments were performed using the perforated patch configuration, except continuous noise recordings in WT and Rh^+/−^ rods using whole-cell patch-clamp. Pipettes were front filled with standard internal solution and back-filled with internal solution containing 30 µM of β-escin (MP Biomedicals, catalog #157941) as perforating agent. For a subset of RBC recordings, 50 µM GTP-γ-S was included in the internal solution, to slowly saturate the RBC transduction cascade (Sampath and Rieke, 2004). Single cell recordings were performed either in voltage or current clamp mode, as previously described (Pahlberg et al., 2018; Bocchero and Pahlberg, 2023). For voltage-clamp recordings, rods were held at -40 mV and RBCs at -60 mV using an Axopatch 200B patch-clamp amplifier (Molecular Devices). Typical seal resistance was ∼1 GΩ, with access resistance of ∼200 MΩ for rods and ∼100 MΩ for RBCs. Membrane potentials were corrected for liquid junction potential (Neher, 1992) previously estimated to be approximately 10 mV for these recording solutions (Campla et al., 2022). Signals were digitized at 10 kHz using a 16-bit A/D converter (ITC18/USB18; HEKA Elektronik) and low-pass filtered at 300 Hz (Frequency Devices 900).

Light-evoked responses were obtained by delivering 20-ms flashes from a blue-green LED (λmax ∼ 505 nm, Thorlabs). Flash strengths ranged from a just measurable response to those that produced a maximal response, generally increasing by a factor of 2. Average dim-flash responses were calculated as the weighted average of all linear range responses, defined as < 25% the maximum response amplitude.

### Physiological recordings of dark noise

Continuous dark noise in rods was isolated from membrane-current fluctuations recorded in darkness as described previously (Baylor et al., 1980; Rieke and Baylor, 1996; Bocchero and Pahlberg, 2023; Stacks et al., 2026). Dark-adapted currents were recorded for 10 s, followed by a 10-s saturating light step to close the light-sensitive current and provide an estimate of instrumental noise. Power spectral densities (PSDs) of the dark and saturated-current recordings were calculated by Fourier transformation using Welch’s method with 50% overlap (MATLAB). The cellular dark-noise PSD was obtained by subtracting the instrumental-noise PSD from the dark-noise PSD (Sampath and Baylor, 2002; Bocchero and Pahlberg, 2023; Stacks et al., 2026).

Dark noise in RBCs was measured using perforated-patch recordings with 50 μM GTP-γ-S included in the intracellular solution. After achieving perforation, 50 s of spontaneous RBC activity was recorded in darkness while intracellular GTP-γ-S remained sufficiently low to preserve normal transduction current. Recordings were repeated after 10–20 min, when intracellular GTP-γ-S had saturated the RBC transduction cascade, providing an estimate of instrumental and residual cellular noise (Sampath and Rieke, 2004). The RBC cellular dark-noise PSD was calculated using the same approach as for rods, by subtracting the PSD measured following GTP-γ-S saturation from the initial dark-noise PSD.

### Electrophysiological data analysis

Data were acquired in MATLAB (R2018b; MathWorks, Natick, MA) using the open-source, Symphony Data Acquisition System (https://symphony-das.github.io). All offline data visualization and analysis were performed using custom MATLAB scripts, Igor Pro, and Metavision (open-source; https://github.com/pahlberglab).

Light sensitivity was estimated from intensity-response relations from each cell using the half-maximal flash strength, which thus is independent of the maximum light amplitude current. Response amplitudes were plotted as a function of flash strength (Φ) and fitted with a Hill equation:

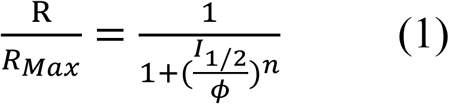

where *R* is the response amplitude, *R*_max_ is the maximal response amplitude, *I*_₁/₂_ is the flash strength producing a half-maximal response, and *n* is the Hill coefficient.

### Signal-to-noise ratio

Signal-to-noise ratio (SNR) in rods and RBCs was determined from response-amplitude distributions generated from repeated dim-flash trials, similar to previously described approaches (Okawa et al., 2010). For each trial, response amplitude was determined from its correlation with the average response across trials. The resulting amplitudes were assembled into histograms and fitted with the sum of three Gaussian distributions using Clampfit 10 (Molecular Devices), representing null events, single-event responses, and double events. The combined Gaussian distributions provided the overall fit to each amplitude histogram. Histograms shown in Figs. 2E and 4E are representative examples from individual cells; the number of trials in these examples does not represent the trial-number distribution for the corresponding genotype.

From the fitted distributions, we extracted D as the mean amplitude of the dark-noise distribution, *A* as the mean amplitude of the single-photon-response distribution, σ_D_ as the standard deviation of the dark-noise distribution, and σ_A_ as the standard deviation of the single-photon-response distribution. SNR was defined as the separation between the noise and single-photon-response distributions normalized by their combined variance.

For rod photoreceptors this becomes:

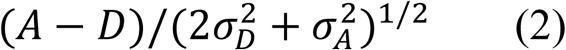

whereas for rod bipolar cells this becomes:

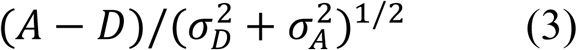

as previously described (Dunn et al., 2006, Okawa et al., 2010). Histogram bin width was determined from the number of trials available for each cell using the same criterion across genotypes and was selected independently of the resulting SNR. Similar numbers of trials were included across genotypes for both rods and RBCs.

### Rod bipolar cell detection threshold

RBC detection threshold was determined from repeated dim-light stimuli ranging from approximately 0.08 to 1 R*/rod. For each flash strength, the fraction of trials producing a detectable RBC response was calculated and plotted as a function of flash strength. The resulting relationship was fitted with a sigmoid function, and the flash strength producing detectable responses in 75% of trials was operationally defined as the detection threshold for that cell.

### Statistical analysis

Unpaired, two-tailed Student’s *t* tests was performed in Excel 2021 to assess continuous noise WT and Rh^+/−^. One-way ANOVA followed by Dunnett’s multiple-comparisons test was used for comparisons among normally distributed groups when individual genotypes were compared with WT. For nonparametric comparisons involving multiple groups, a Kruskal–Wallis test followed by Dunn’s multiple-comparisons test was used. Statistical significance was defined as * = p < 0.05, ** = p < 0.01, *** = p < 0.001. Data are reported as the mean ± SEM and number of cells (*N*), unless otherwise noted. All electrophysiology figures were made using Igor 8.0 from WaveMetrics.

### Data Availability

Data supporting the findings of this study and custom analysis code are available from the corresponding author upon reasonable request.

## Results

### Dark continuous noise in rod photoreceptors

*Continuous* dark noise in rod photoreceptors consists of random fluctuations in outer-segment current, and the consensus for decades has been that it is produced by spontaneous activity within the phototransduction cascade, more specifically the dark basal activity of PDE6 (Baylor et al., 1980; Rieke and Baylor, 1996; Lamb, 1987; Bocchero and Pahlberg, 2023; Stacks et al., 2026). Recently, an alternative model proposed that a substantial component of continuous noise arises from the visual pigment itself, rather than from spontaneous PDE6 activity (Chai et al., 2024). We therefore further examined the relationship between rhodopsin concentration and continuous noise by comparing WT and Rh^+/−^ rods in a normal C57BL/6J background (Fig. 1).

**Figure 1.**
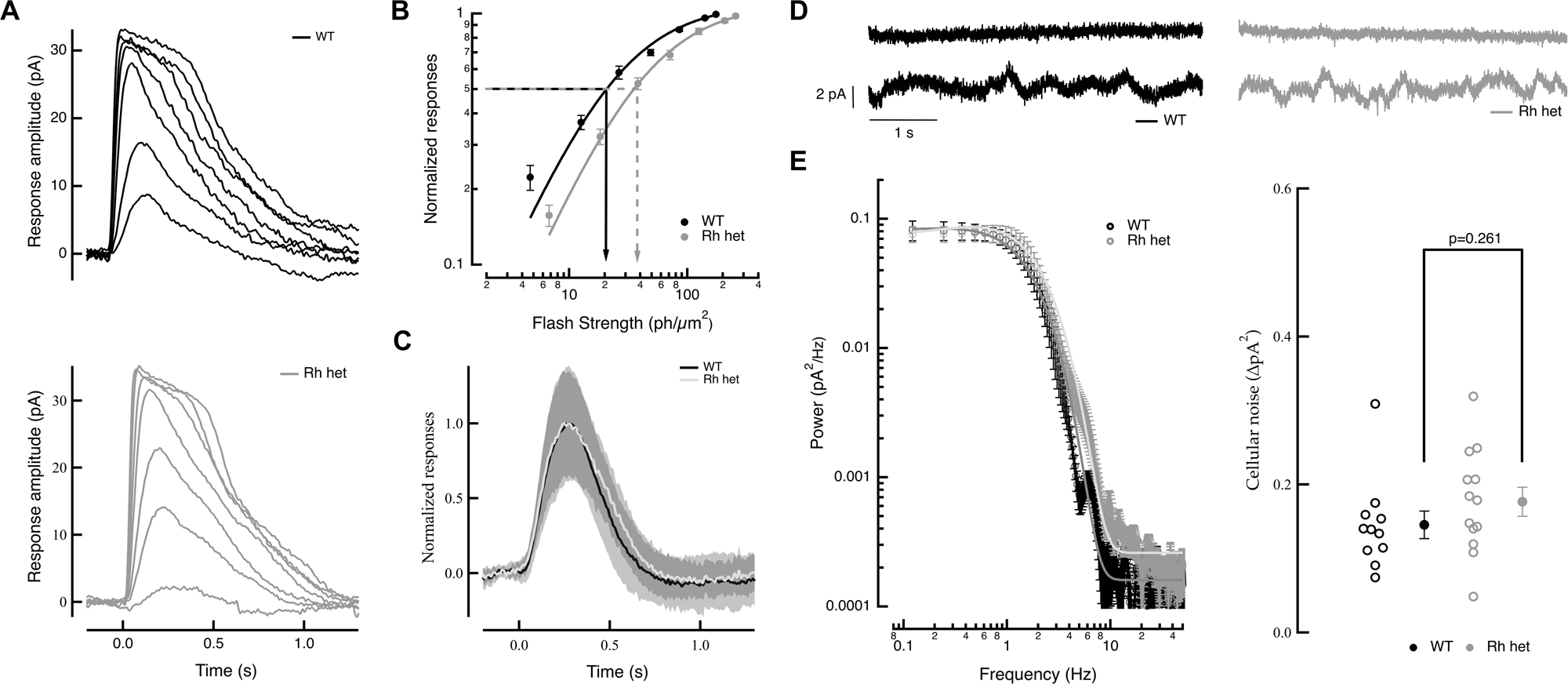
Continuous noise is independent of rhodopsin concentration. **A)** Representative light-evoked flash families from WT (black) and Rh^+/-^(gray) rods are similar. Patch-clamp recordings were made in whole-cell mode [membrane potential (*V*_m_) = −40 mV]. Twenty millisecond light flashes given at time 0 yielded flash strengths ranging from 0.3 to 174.5 R*/rod for WT and 0.5 to 257 R*/rod for Rh^+/-^, increasing by a factor of 2. Recordings are representative of data collected from several experiments. **B)** Intensity–response plot (mean ± SEM) of WT rods (black; *n* = 14) and Rh^+/−^ rods (grey; *n* = 30). Rh^+/-^ are ∼ 2-fold less sensitive than WT rods. The sensitivity (*I*_1/2_) was derived from the best fitting Hill equation, *I*_1/2_ = 22.2 ± 1.7 and 42.5 ± 2 R*/rod, respectively. **C)** Average normalized dim-flash responses recorded with the patch-clamp technique from WT (black) and Rh ^+/−^ (gray) rods. Responses were derived from 5 WT and 4 Rh ^+/−^ rods (<25% of maximum response amplitude), and the time-to-peak values were 246 ± 15.1 and 237.5 ± 18.8 ms, respectively (mean ± SEM).**D)** Representative recordings of membrane currents from a WT littermate rod (black) and a Rh ^+/-^ rod (gray). Whole-cell patch-clamp recordings were done in complete darkness, followed by a saturating light flash (8725 R*/rod/s) to close all of the CNG channels. **E)** Continuous noise power spectra from WT (black, *n* = 11) and *Rh*^+/−^ (gray, *n* = 13) rods (mean ± SEM). Spectra were constructed by fast Fourier transformation of the cellular dark noise, derived by subtracting the instrumental noise (saturated response) from dark noise recordings. F) Cumulative power of dark noise between 0.1 and 10 Hz (mean ± SEM). No statistically significant (*p* = 0.26095; Student’s t-test) difference could be found in dark noise between WT (black) and *Rh* ^+/-^ (gray) rods.

As we previously showed (Bocchero and Pahlberg, 2023), both genotypes generated robust light-evoked responses with similar response kinetics (Fig. 1A). As expected from their reduced pigment concentration, Rh^+/−^ rods were approximately twofold less sensitive than WT rods (Fig. 1B), while normalized dim-flash responses showed similar kinetics between genotypes (Fig. 1C). The similar kinetics are consistent with our previous measurements in Rh^+/−^ mice on the same C57BL/6J background (Bocchero and Pahlberg, 2023). Accelerated Rh^+/−^ response kinetics have been reported in other preparations (Calvert et al., 2001), and may depend on genetic background and associated differences in rod outer-segment dimensions (Okawa and Sampath, 2025). Thus, reducing rhodopsin concentration reduced rod sensitivity without substantially altering response kinetics under the conditions used here.

We next measured continuous noise as previously described (Fig. 1D). WT and Rh^+/−^ rods showed closely overlapping noise spectra (Fig. 1E), and cumulative noise power between 0.1 and 10 Hz was not significantly different between genotypes (Fig. 1F). Thus, reducing rhodopsin concentration sufficiently to produce an approximately twofold decrease in light sensitivity, and previously shown to produce a corresponding reduction in discrete noise events (Bocchero and Pahlberg, 2023), did not produce a reduction in continuous dark noise. Together with our previous findings that continuous noise is independent of transducin concentration, increases following reduction of PDE6 expression (Bocchero and Pahlberg, 2023), and decreases following deletion of the PDE6 modulator GARP2 (Stacks et al., 2026), these results strongly support basal PDE6 activity as a major determinant of continuous dark noise.

### Manipulation of phototransduction noise alters signal fidelity in rods

To determine how discrete and continuous dark noise limit signal fidelity at the photoreceptor level, we compared the ability of rods with selectively altered noise components to distinguish photon-driven responses from intrinsic noise (Fig. 2). First, flash families and intensity-response relationships were obtained to characterize normal cellular responses and sensitivity (Fig. 2A and B). Next, we delivered repeated dim flashes near the threshold for photon detection. Because photon absorption follows Poisson statistics, repeated weak flashes produced a mixture of light-evoked responses and null events in which no photon response was detected (Fig. 2C). From these data we constructed response-amplitude histograms and fitted them with Gaussian distributions representing null-and photon-driven events to quantify their separation and derive a measure for the signal-to-noise ratio (SNR; Fig. 2D; see Okawa et al., 2010).

**Figure 2.**
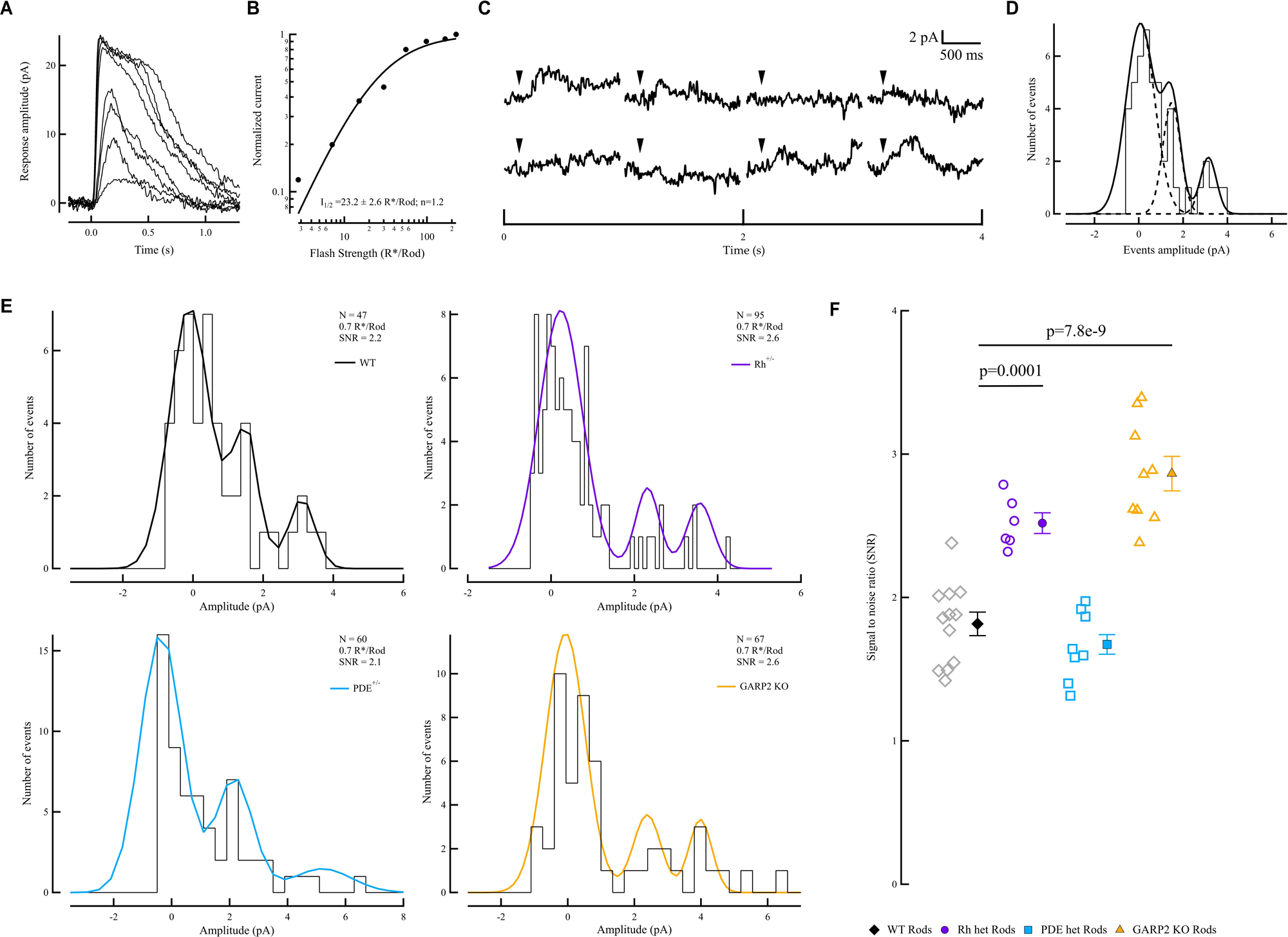
Signal-to-noise ratio in rods. A) Representative light-evoked flash family (FF) from a WT rod under perforated-patch conditions. B) Intensity-response curve from a representative WT rod to assess sensitivity (I_1/2_). C) Dim-flash responses from a WT rod recorded with the perforated- patch-clamp technique. Twenty-millisecond light flashes are indicated by arrows. D) Schematic amplitude histogram of dim-flash and null events. The bold black line is the best fit of the histogram profile (three Gaussian distributions, dashed lines) used to derive the signal-to-noise ratio (SNR). E) Representative event distributions from individual WT (black), Rh^+/−^ (magenta), PDE6^+/−^ (blue), and GARP2^−/−^ (yellow) rods. Trial counts shown for representative histograms refer to the individual cells displayed and do not represent genotype-level trial numbers. F) SNR in WT (N=12), Rh^+/−^ (N=6), PDE6^+/−^ (N=8), and GARP2^−/−^ (N=9) rods (mean ± SEM). SNR was significantly increased in Rh^+/−^ (p = 0.0001) and GARP2^−/−^ (p < 0.0001) rods compared with WT.

Response-amplitude distributions showed clear genotype-dependent differences in the separation of photon-driven responses from intrinsic noise among WT, Rh^+/−^, PDE6^+/−^, and GARP2^−/−^ rods (Fig. 2E). Quantification revealed a significantly increased SNR in GARP2^−/−^ rods in which continuous noise is reduced, and surprisingly also in Rh^+/−^ rods, in which discrete noise is reduced, compared to WT (*p* < 0.001; Fig. 2F). Thus, reducing either form of dark noise increased rod SNR, indicating that both discrete and continuous noise contribute to limiting signal fidelity in individual rods.

### Changes in continuous rod noise are transmitted to rod bipolar cells

RBCs receive convergent synaptic input from multiple rods. We therefore asked whether differences in continuous noise remain detectable following convergence at the rod-to-RBC synapse. Membrane-current fluctuations were again recorded in darkness using perforated-patch clamp (Fig. 3A). GTP-γ-S, a poorly hydrolysable analog of GTP (Lamb and Matthews, 1988), was used to close TRPM1 channels, providing an estimate of the current fluctuations that remained after elimination of the RBC transduction-channel component. Subtraction of this residual component from the dark recording was used to estimate cellular RBC noise, which was then analyzed in the frequency domain (Fig. 3B). As in rod photoreceptors, RBC noise power spectra differed between genotypes. WT and Rh^+/−^ mice showed largely overlapping noise spectra, whereas the increase in continuous rod noise associated with reduced PDE6 expression remained apparent at the level of the RBC (Fig. 3C), although it did not reach significant levels. Conversely, the reduction in continuous rod noise associated with GARP2 deletion was preserved in the RBCs, with cumulative noise power between 0.1 and 10 Hz significantly lower in GARP2−/− RBCs than in WT RBCs (p < 0.05; Fig. 3D). Thus, differences in continuous rod noise remain detectable following synaptic transmission and convergence onto RBCs, demonstrating that presynaptic phototransduction noise contributes to the noise in RBCs.

**Figure 3.**
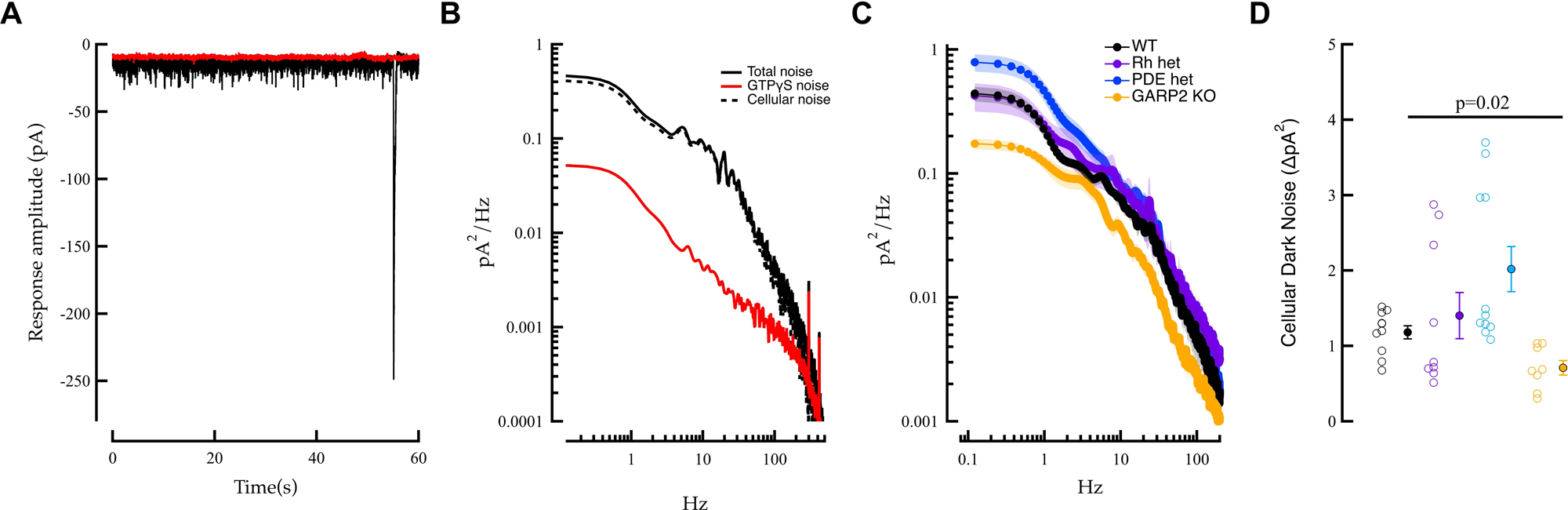
Dark noise in RBCs. A) Representative membrane-current recordings from RBCs using the perforated-patch technique in darkness (black trace) and after GTP-γ-S saturation of the transduction cascade (red trace). B) Power spectra of dark noise and residual noise from a representative RBC. The dotted gray power spectrum represents total cellular dark noise obtained by subtracting the red trace from the black trace. C) Power spectra of cellular noise in WT (black; N=14), PDE6^+/−^ (N=9), and GARP2^−/−^ (N=8) RBCs. D) Cumulative dark-noise power between 0.1 and 10 Hz (mean ± SEM; units pA²). Dark noise was significantly reduced in GARP2^−/−^ RBCs (p = 0.020; Kruskal-Wallis test).

### Continuous rod noise controls signal-to-noise ratio in rod bipolar cells

Because RBCs receive convergent input from multiple rods, they integrate sparse photon-driven signals together with intrinsic noise generated across the presynaptic rod population (Baylor et al., 1984; van Rossum and Smith, 1998). Signal transmission at the rod-to-RBC synapse is strongly nonlinear, providing a mechanism to suppress weak rod inputs and continuous noise while preferentially transmitting larger single-photon responses (SPRs) (Field and Rieke, 2002; Robson et al., 2004). We therefore asked how differences in rod signal and noise distributions affect signal fidelity following transmission through this nonlinear synapse. As for the rod recordings, we first measured families of light-evoked responses and constructed intensity-response relationships for individual RBCs (Fig. 4A, B). Repeated dim flashes were then delivered to generate populations of light-evoked responses and null trials (Fig. 4C), from which response-amplitude histograms were constructed and fitted with Gaussian distributions as described for rods (Fig. 4D).

**Figure 4.**
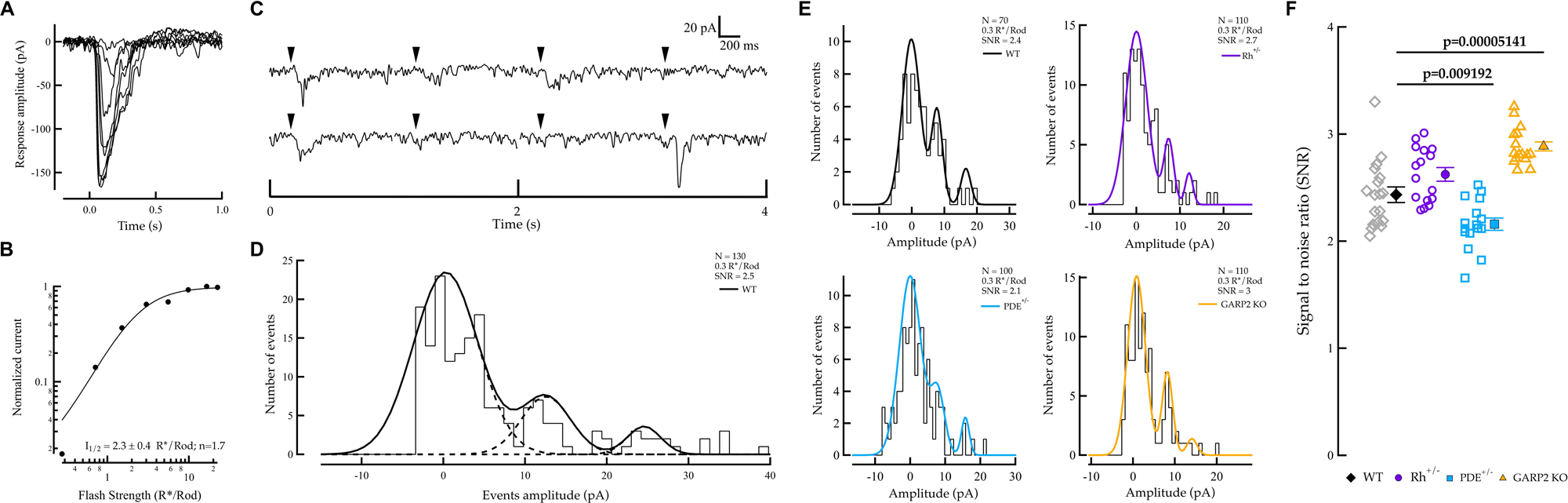
Signal-to-Noise ratio in RBCs. **A)** Representative light-evoked flash family from a WT RBC. **B)** Intensity-Response curve from a WT RBC to assess light sensitivity (I_1/2_). **C)** Dim flash responses from a WT RBC recorded with the perforated patch-clamp technique. Twenty millisecond light flashes indicated by arrows. **D)** Schematic amplitude histogram of dim flash and null events. Bold black line is the best fit of the histogram profile (three gaussian distributions dashed lines) to derive the signal-to-noise ratio (SNR). **E)** Representative event distributions for WT (black) Rh ^+/-^ (magenta) PDE ^+/-^ (blue) and GARP2 ^-/-^ (yellow) RBCs. **F**) SNR in WT (N=19), Rh^+/-^(N=18), PDE^+/-^ (N=17) and GARP2^-/-^ (N=19) RBCs (mean ± SEM). SNR was significantly decreased in PDE6+/− (p = 0.009) and increased in GARP2−/− (p < 0.0001) RBCs compared with WT.

Response-amplitude distributions again showed genotype-dependent differences in the separation of light-evoked responses from intrinsic RBC noise (Fig. 4E). In WT mice, SNR increased significantly from rods to RBCs (*p* < 0.001; Fig. 4F), consistent with the established role of the rod-to-RBC nonlinearity in suppressing continuous noise while preferentially transmitting larger SPRs (Field and Rieke, 2002; Robson et al., 2004). In contrast, SNR in Rh+/− RBCs was not significantly different from WT RBCs. Thus, although reducing discrete noise increased SNR within Rh^+/−^ rods (Fig. 2F), this advantage was not maintained following transmission to RBCs.

Despite the increased continuous noise in PDE6^+/−^ rods, SNR increased following transmission to RBCs, indicating that the rod-to-RBC nonlinearity still improved signal separation. However, SNR in PDE6^+/−^ RBCs remained significantly lower than in WT RBCs (*p* < 0.01; Fig. 4F). In contrast, GARP2^−/−^ rods already exhibited increased SNR associated with reduced continuous noise, and no further increase in SNR was observed following transmission to RBCs. Nevertheless, SNR in GARP2^−/−^ RBCs remained significantly higher than in WT RBCs (*p* < 0.001; Fig. 4F).

### Nonlinear processing at the rod-to-RBC synapse determines the impact of continuous noise

Previous work has demonstrated that the threshold-like nonlinearity in RBCs is created by saturation within the postsynaptic G-protein signaling cascade (Sampath and Rieke, 2004). Our results indicate that the effectiveness of nonlinear filtering is altered depending on the signal and noise distributions of the presynaptic rod population (Fig. 4). We therefore tested whether reducing the rod-to-RBC nonlinearity alters the impact of continuous rod noise on postsynaptic SNR.

The nonlinearity was manipulated pharmacologically using the mGluR6 antagonist LY-341495. Representative response families were obtained before and during LY-341495 application (Fig. 5A). Intensity-response relationships demonstrated a significant reduction in response nonlinearity following LY-341495 application (Fig. 5B), accompanied by a change in RBC holding current (Fig. 5C). This reduction of the nonlinearity altered the genotype-dependent differences in RBC SNR (Fig. 5D). PDE6^+/−^ RBCs continued to exhibit significantly lower SNR than WT RBCs (p = 0.018). In contrast, GARP2^−/−^ RBCs, which exhibited significantly higher SNR than WT under control conditions (Fig. 4F), were no longer significantly different from WT following LY-341495 treatment (Fig. 5D). Thus, reducing the nonlinearity eliminated the SNR advantage associated with reduced continuous noise in GARP2^−/−^ retina. Together, these findings indicate that the effect of altered continuous rod noise on postsynaptic SNR depends on nonlinear processing at the rod-to-RBC synapse.

**Figure 5.**
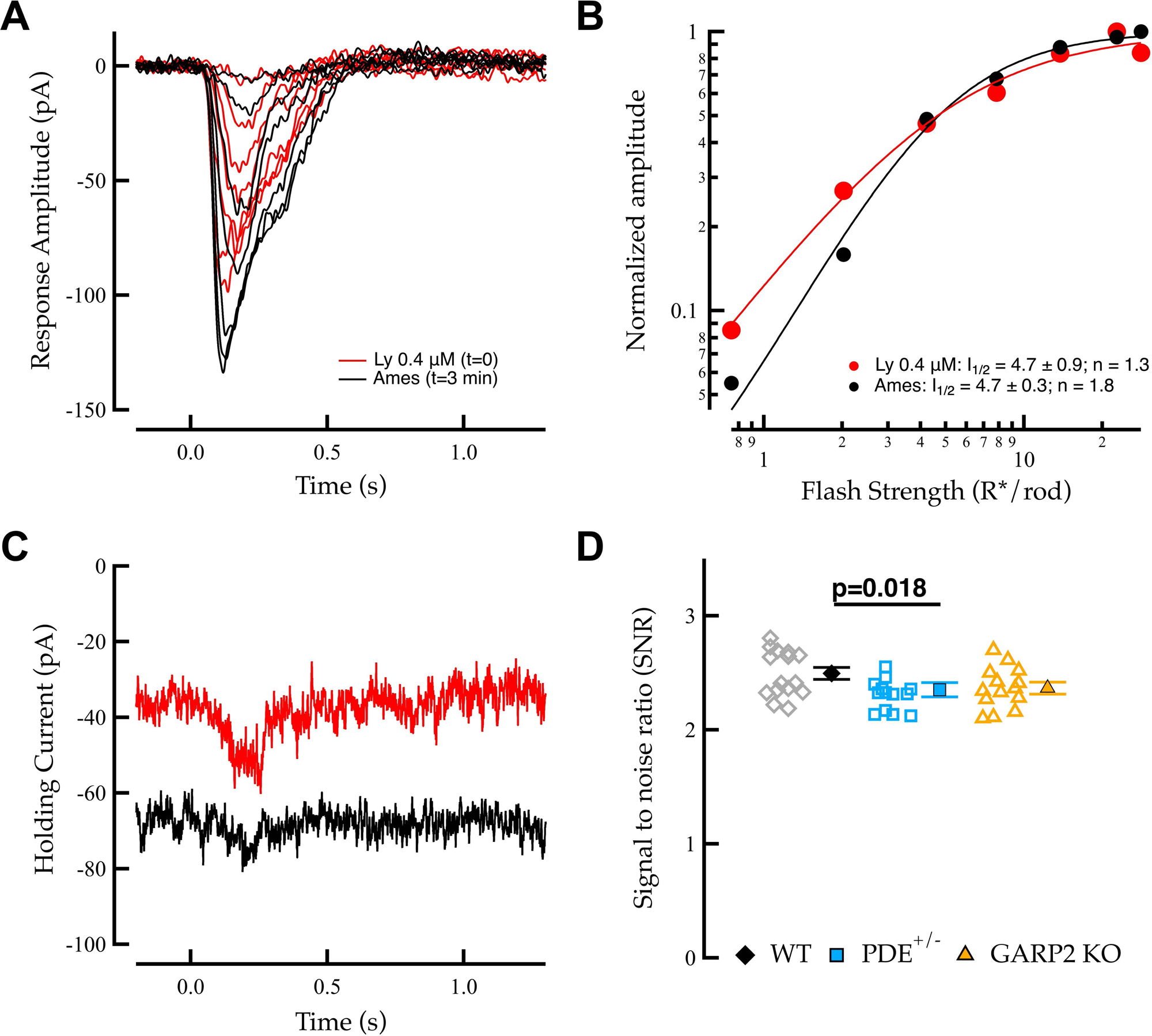
Reduction of nonlinearity in RBCs affects SNR. A) Representative light-evoked flash families from an RBC in Ames medium (black) and during LY-341495 application (red). B) Intensity-response curves before (black) and during LY-341495 application (red) used to assess sensitivity (I_1/2_) and nonlinearity (n). C) Change in RBC holding current during LY-341495 application. D) SNR in RBCs during LY-341495 application for WT (N=15), PDE6^+/−^ (N=14), and GARP2^−/−^ (N=14). SNR was significantly lower in PDE6+/− RBCs than in WT (p = 0.018), whereas GARP2^−/−^ RBCs did not differ significantly from WT (p = 0.125).

### Reduced continuous noise improves detection of near-threshold signals by rod bipolar cells

Finally, we asked whether changes in continuous noise and SNR altered the ability of RBCs to detect weak light signals near detection threshold. RBC responses were recorded using 20-ms flashes ranging from approximately 0.08 to 1 R*/rod (Fig. 6A). For each flash strength, detection performance was quantified as the fraction of trials producing a detectable RBC response. The fraction detected increased with flash strength in all genotypes (Fig. 6B), and the flash strength producing detectable responses in 75% of trials was used to define the operational RBC detection threshold.

**Figure 6.**
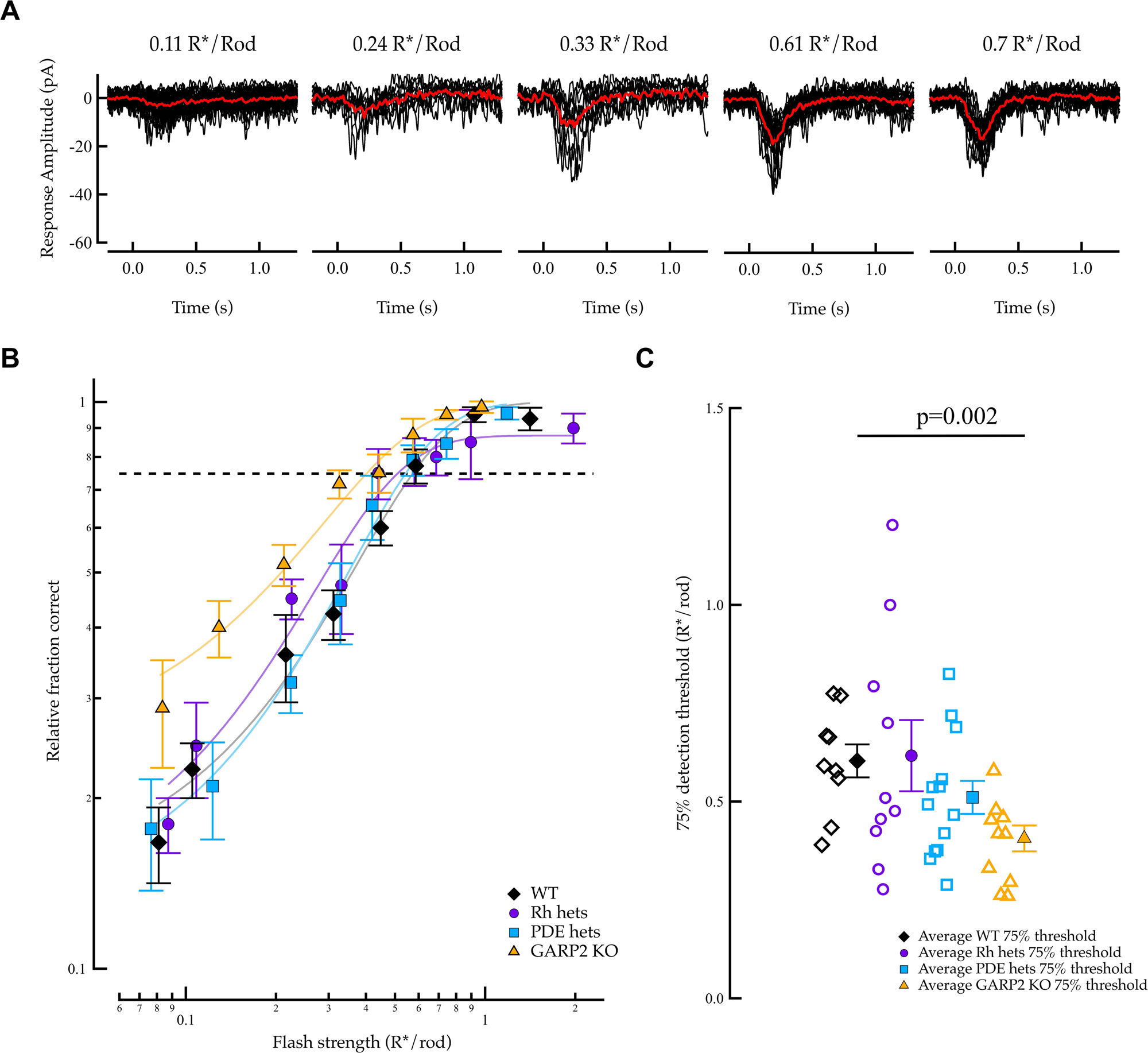
Detection threshold in RBCs. A) Representative recordings of near-threshold responses using the perforated-patch-clamp technique. Arrows indicate 20-ms light flashes. Flash intensities ranged from approximately 0.08 to 1.1 R*/rod. B) Fraction of trials producing a detectable RBC response (mean ± SEM) as a function of flash strength for WT (N=9), Rh+/− (N=10), PDE6^+/−^ (N=13), and GARP2^−/−^ (N=10) RBCs. The dashed line indicates the 75% detection criterion. C) Average RBC detection threshold. Detection threshold was significantly lower in GARP2^−/−^ RBCs than in WT (p = 0.002; Kruskal-Wallis test).

GARP2^−/−^ RBCs, which exhibited reduced postsynaptic dark noise and increased SNR (Figs. 3, 4), also showed a significantly lower detection threshold than WT RBCs (*p* < 0.01; Fig. 6C). This indicates that the improved SNR associated with reduced continuous rod noise was preserved through the rod-to-RBC synapse and accompanied by improved detection of near-threshold light responses. Thus, the level of continuous phototransduction noise in rods contributes significantly to setting the detection threshold of the primary rod pathway.

## Discussion

The performance of the visual system near absolute threshold is ultimately constrained by intrinsic noise generated within the retina, which limits the ability to distinguish signals produced by photon absorption from spontaneous activity in darkness (Barlow, 1956; Aho et al., 1987, 1988). Here, we used genetically modified mouse models to investigate the molecular mechanism of continuous noise and to determine how intrinsic phototransduction noise influences signal transmission at the first visual synapse. Reducing rhodopsin concentration did not reduce continuous dark noise power, further supporting basal PDE6 activity as a major source of continuous noise under our experimental conditions (Fig. 1). Reducing either discrete noise or continuous noise increased SNR within individual photoreceptors (Fig. 2), demonstrating how both forms of intrinsic noise can limit signal fidelity at the rod level. However, only changes in continuous noise had a strong effect on signal processing following transmission to rod bipolar cells. Reduced continuous noise decreased RBC dark noise, increased RBC SNR, and lowered detection threshold, whereas increased continuous noise resulted in reduced postsynaptic SNR (Figs. 3, 4, 6). Moreover, the rod-to-RBC nonlinearity substantially increased SNR under WT conditions but was less effective when presynaptic continuous noise was experimentally altered, and pharmacological reduction of this nonlinearity changed the genotype-dependent differences in RBC SNR (Figs. 4, 5). Taken together, these results demonstrate that continuous phototransduction noise and nonlinear processing at the rod-to-RBC synapse interact to determine the fidelity and threshold of dim-light signal transmission.

### Molecular origin of continuous dark noise

Continuous rod noise has traditionally been thought to arise from spontaneous activity downstream of rhodopsin, with PDE6 considered a likely source (Baylor et al., 1980; Lamb, 1987; Rieke and Baylor, 1996). Our previous functional experiments on genetically modified mice supported this interpretation, as continuous noise was independent of rhodopsin and transducin concentration, increased following reduction of PDE6 expression (Bocchero and Pahlberg, 2023), and decreased following deletion of the PDE6 regulator GARP2 (Stacks et al., 2026).

Chai et al. (2024) recently proposed instead that a substantial component of continuous noise originates from a metastable state of visual pigment. Their principal experiments, however, used REY-rhodopsin, a genetically modified pigment with severely impaired transducin activation, on a GCAPs^−/−^ background that itself substantially alters continuous-noise properties. REY-Rho;GCAPs^−/−^ rods were also ∼7,400-fold less light sensitive than WT-Rho;GCAPs^−/−^ rods, making direct comparison with rods expressing native pigment difficult. Our Rh^+/−^ experiments provide a complementary test in rods containing native mouse rhodopsin. Reducing rhodopsin concentration by approximately one-half reduced light sensitivity but did not reduce continuous-noise power (Fig. 1). If continuous noise arose predominantly from spontaneous activity of rhodopsin, its magnitude would be expected to scale with pigment concentration, as occurs for discrete noise (Bocchero and Pahlberg, 2023). Together with the increased continuous noise following reduction of PDE6 expression (Bocchero and Pahlberg, 2023) and decreased continuous noise following GARP2 deletion (Stacks et al., 2026), these results support basal PDE6 activity as a major determinant of continuous dark noise in WT rods.

An important and somewhat unexpected finding of the present study is that reducing *either* form of intrinsic noise can improve signal fidelity within individual photoreceptors. GARP2^−/−^ rods, in which continuous noise is reduced, showed increased SNR as expected from the narrower background-noise distribution. Surprisingly, SNR was also increased in Rh^+/−^ rods, in which the frequency of discrete events is reduced (Fig. 2F). Because these spontaneous events closely resemble genuine single-photon responses in amplitude and kinetics (Baylor et al., 1980; Rieke and Baylor, 1998; Bocchero and Pahlberg, 2023), reducing their frequency would not be expected to improve discrimination between an individual discrete event and an SPR. The increased SNR in Rh^+/−^ rods therefore suggests that reducing the frequency of spontaneous photon-like events alters the overall response distribution, although the mechanism by which this is reflected in the measured SNR remains unclear. Thus, although discrete and continuous noise arise through distinct molecular mechanisms, both contribute to limiting signal fidelity within individual rods, consistent with the longstanding view that intrinsic photoreceptor noise constrains visual sensitivity (Barlow, 1956; Donner, 1992; Rieke and Baylor, 1998; Pahlberg and Sampath, 2011).

### Rod continuous noise is a major determinant of signal processing in the primary rod pathway

The consequences of rod noise become particularly important after convergence onto RBCs, where signals and noise from multiple photoreceptors must be integrated (Baylor et al., 1984). Our results indicate that their consequences differ following transmission through the rod-to-RBC synapse. Notably, changes in continuous rod noise remained detectable at the level of the RBC, where GARP2^−/−^ RBCs exhibited significantly lower dark-noise power than WT RBCs (Fig. 3), consistent with the reduction in continuous noise previously observed in GARP2^−/−^ rods (Stacks et al., 2026). In WT retina, SNR increased significantly from rods to RBCs (Fig. 4F), consistent with the role of the rod-to-RBC nonlinearity in filtering rod noise while preserving single-photon responses (Field and Rieke, 2002; Sampath and Rieke, 2004). Although reducing discrete noise increased SNR in Rh^+/−^ rods, this improvement was not maintained at the RBC level. Because continuous noise was similar in WT and Rh^+/−^ rods (Fig. 1), reducing the frequency of discrete events could be expected to have relatively little effect on the continuously present noise distribution seen by the rod-to-RBC nonlinearity.

Manipulations of continuous noise, however, produced a different pattern. The comparable SNR of WT and Rh^+/−^ RBCs suggests that continuous noise has a greater influence than discrete-event frequency on the increase in SNR across the synapse. Indeed, increased continuous noise in PDE6^+/−^ rods limited the gain in SNR following transmission to RBCs compared to WT. In contrast, GARP2^−/−^ rods that already exhibited high SNR because of reduced continuous noise, showed little additional increase following transmission to RBCs. Nevertheless, postsynaptic SNR remained significantly higher in GARP2^−/−^ than WT (Fig. 4F).

The detection-threshold measurements further highlight the functional importance of continuous noise in the primary rod pathway. Continuous noise is constantly present across the population of rods converging onto an RBC and therefore contributes to the background against which sparse photon responses must be detected (Baylor et al., 1980; van Rossum and Smith, 1998). The higher SNR present in GARP2^−/−^ rods was maintained following transmission to RBCs and was associated with a lower threshold for detecting weak light stimuli (Fig. 6C). Thus, reducing continuous phototransduction noise improves both SNR and the ability of RBCs to discriminate weak light responses from background noise, improving detection of near-threshold light responses.

### Nonlinear thresholding of rod input

The effect of continuous rod noise on downstream signaling in the primary rod pathway is shaped by the nonlinear transformation at the rod-to-RBC synapse, and previous work demonstrated that saturation within the postsynaptic signaling cascade creates the threshold-like nonlinearity (Field and Rieke, 2002; Sampath and Rieke, 2004). This mechanism is particularly advantageous in the convergent rod pathway, where sparse photon-driven signals must be preserved while noise contributed by multiple presynaptic rods is suppressed (van Rossum and Smith, 1998; Pahlberg and Sampath, 2011).

The effectiveness of the nonlinearity depends on the relative distributions of signal and noise arriving from the converging rod population and on the position of the threshold separating them (van Rossum and Smith, 1998; Field and Rieke, 2002; Taylor and Smith, 2004). Our results suggest that this threshold is well positioned for the normal signal and noise distributions of WT rods, where SNR increased substantially following transmission to RBCs. When continuous noise was increased in PDE6^+/−^ rods, SNR was still improved but remained lower than WT levels, suggesting that the normal threshold was less effective at separating photon responses from the broadened noise distribution. In contrast, GARP2^−/−^ rods already exhibited greater separation between signal and noise because of reduced continuous noise, and little additional improvement in SNR occurred at the RBC level. Nevertheless, SNR in GARP2^−/−^ RBCs remained significantly higher than in WT, consistent with the narrower noise distribution lying predominantly below the nonlinear threshold. Thus, these findings indicate that as previously hypothesized, the effectiveness of nonlinear filtering depends strongly on the signal and noise distributions presented to the synapse (Field and Rieke, 2002; Sampath and Rieke, 2004; Pahlberg and Sampath, 2011).

The LY-341495 experiments further support this conclusion. Reducing the nonlinearity altered the differences in RBCs observed under control conditions (Fig. 5). In PDE6^+/−^ RBCs, the SNR remained significantly lower compared to WT, suggesting that elevated continuous noise in the presynaptic rods continued to limit signal detection even when the nonlinearity was reduced. In contrast, reducing the nonlinearity abolished the SNR advantage of GARP2^−/−^ RBCs compared to WT. The normal rod-to-RBC nonlinearity appears well matched to the signal and noise statistics of WT rods but does not fully adjust when presynaptic noise distributions are altered. Increased continuous noise limits the efficiency of nonlinear filtering, whereas reduced continuous noise leaves less noise to be removed by nonlinear filtering at the synapse. Thus, postsynaptic SNR depends not only on the level of rod noise, but also on the position of the nonlinear threshold relative to the presynaptic signal and noise distributions.

Together, our findings link the molecular origin of phototransduction noise to signal processing in the primary rod pathway. Basal PDE6 activity is a major determinant of continuous rod noise, which shapes the signaling from rods to RBCs. Nonlinear processing at the rod-to-RBC synapse further determines how this noise is separated from photon-driven signals and how faithfully near-threshold responses are transmitted to the downstream circuitry. Overall, sensitivity near visual threshold in the primary rod pathway depends on both the coordinated control of phototransduction noise in rods and its nonlinear filtering at the first visual synapse.

## Author contributions

U.B. and J.P. designed research; U.B. and J.P. performed research; U.B. and J.P. analyzed data; U.B. and J.P. edited the paper; U.B. and J.P. wrote the paper.

This work was supported by the Intramural Research Programs of NEI/NIDCR/NINDS at the National Institutes of Health Grants 100000072 and 100000065 (JP). The contributions of the NIH author(s) were made as part of their official duties as NIH federal employees, are in compliance with agency policy requirements, and are considered Works of the United States Government. However, the findings and conclusions presented in this paper are those of the author(s) and do not necessarily reflect the views of the NIH or the U.S. Department of Health and Human Services.

## Acknowledgements

We thank Dr. Steven J. Pittler for generously providing the GARP2^−/−^ mice, Jeannie Chen for the Rh^+/−^ mice, and Dr. Nikolai O. Artemyev for the PDE6^+/−^ mice. We thank Dr. Alapakkam Sampath for comments and suggestions on the manuscript.

The authors declare no competing interests.

